# Reduced language lateralization in autism and the broader autism phenotype as assessed with robust individual-subjects analyses

**DOI:** 10.1101/2020.02.10.942698

**Authors:** Olessia Jouravlev, Alexander J.E. Kell, Zachary Mineroff, A.J. Haskins, Dima Ayyash, Nancy Kanwisher, Evelina Fedorenko

## Abstract

One of the few replicated functional brain differences between individuals with autism spectrum disorders (ASD) and neurotypical (NT) controls is reduced language lateralization. However, most prior reports relied on comparisons of group-level activation maps or functional markers that had not been validated at the individual-subject level, and/or used tasks that do not isolate language processing from other cognitive processes, complicating interpretation. Furthermore, few prior studies have examined functional responses in other functional networks, as needed to determine the selectivity of the effect. Using fMRI, we compared language lateralization between 28 ASD participants and carefully pairwise-matched controls, with the language regions defined individually with a well-validated language localizer. ASD participants showed less lateralized responses due to stronger right hemisphere activations. Further, this effect did not stem from a ubiquitous reduction in lateralization across the brain: ASD participants did not differ from controls in the lateralization of two other large-scale networks—the Theory of Mind network and the Multiple Demand network. Finally, in an exploratory study, we tested whether reduced language lateralization may also be present in NT individuals with high autistic trait load. Indeed, autistic trait load in a large set of NT participants (n=189) was associated with less lateralized language activations. These results suggest that reduced language lateralization is a robust and spatially selective neural marker of autism, present in individuals with ASD, but also in NT individuals with higher genetic liability for ASD, in line with a continuum model of underlying genetic risk.

## 1. Introduction

Many differences in brain structure and function once thought to be hallmarks of autism (Just et al., 2004; Hughes, 2009; Pierce et al., 2001; Gallagher et al., 2000) have turned out to be unreliable or artifactual (Hadjikhani et al., 2007; Deen & Pelphrey, 2012; Dufour et al., 2013; Koldewyn et al., 2014; He et al., 2020). One finding has withstood the test of time and replication: reduced lateralization during speech/language processing (Herbert et al., 2002; Kleinhans et al., 2008; Knaus et al., 2008; 2010; Eyler et al., 2012; see Lindell & Hudry, 2013, for a review, and Herringshaw et al., 2016, for a meta-analysis). Phenotypically, deficits in language and communication are a core feature of autism spectrum disorders (ASD) (Tager-Flusberg et al., 2013; Lord et al., 2012; Wilkinson, 1998; Volkmar et al., 2004). Reduced language lateralization might therefore be a neural marker of communicative impairment in ASD.

Although reduced language lateralization appears to be consistent across paradigms and studies, the majority of prior work (cf. Kleinhans et al., 2008) has relied on comparisons of group-level activation maps. Because individuals vary in the precise locations of macro- and micro-anatomical areas (Amunts et al., 1999; Juch et al., 2005; Tomaiuolo et al., 1999), functional activations do not align perfectly, especially in the higher-order association cortex (Fischl et al., 2008; Frost & Goebel, 2012; Vázquez-Rodríguez et al., 2019). This variability may further be greater in populations with neurodevelopmental disorders like autism (Müller et al., 2003). Apparent reduction in activity in some brain areas in ASD at the group level—which would translate in reduced lateralization if these were left hemisphere (LH) areas—may therefore simply reflect higher variability in the locations of the relevant functional regions.

Furthermore, the nature and scope of lateralization reduction in ASD remain poorly understood. ***First***, the cause of the reduced lateralization is debated. Some report decreased activity in the LH (Harris et al., 2006; Eyler et al., 2012; Müller et al., 1999); others—increased activity in the RH (Anderson et al., 2010; Takeuchi et al., 2004; Knaus et al., 2008; Tesink et al., 2009; Wang et al., 2006); yet others find both (Kleinhans et al., 2008; Boddaert et al., 2003; Redcay & Courchesne, 2008). Given that LH and RH language regions plausibly contribute differently to language processing (Lindell, 2006; Mitchell & Crow, 2005), understanding which hemisphere drives the lateralization differences is critical for interpretation. ***Second***, many studies use language tasks that are not designed to isolate particular cognitive processes. For example, verbal-fluency tasks, where participants are presented with a cue (letter, category name, etc.) and are asked to generate as many associated words as possible (Kenworthy et al., 2013; Kleinhans et al., 2008) certainly engage linguistic resources—spanning several aspects of language (Bradshaw et al., 2017)—but also have an executive component (Thompson-Schill et al., 1997). As a result, these tasks may activate multiple functionally distinct brain networks, complicating the interpretation of the lateralization differences. And ***third***, most studies examine functional responses during a single task making it impossible to determine whether reduced lateralization is specific to that task or the brain network(s) that the task targets, or whether it instead stems from an across-the-brain reduction in lateralization (Dawson, 1983; Fein et al., 1984; Cardinale et al., 2013).

To illuminate the nature and scope of reduced language lateralization in autism, in **Study 1**, we compared individuals with an ASD diagnosis and pairwise-matched controls using a well-validated language “localizer” (Fedorenko et al., 2010) to identify language-responsive regions in each brain individually. This task selectively targets the fronto-temporal language network (Fedorenko et al., 2011; Blank et al., 2014). We additionally included two other localizers each targeting a network implicated in higher-level cognition—the network that supports social cognition, including Theory of Mind (Saxe & Kanwisher, 2003) and the Multiple Demand network (Duncan, 2010, 2013)—to test whether lateralization reduction is restricted to the language network.

Moreover, the emerging genetic picture of autism is complex, with numerous genes implicated and genetically-mediated overlaps with autistic-like traits present in the general population (Huguet et al., 2013; Miles, 2011; Talkowski et al., 2014). If reduced language lateralization is a true endophenotype of autism, variability in lateralization among neurotypical (NT) individuals should relate to autistic trait load (Baron-Cohen et al., 2011; Lai et al., 2015). In a more exploratory **Study 2**, we examined variability in language lateralization in a large NT population (n=189) as a function of autistic trait load.

## 2. Methods

### 2.1. Participants

Participants gave informed consent in accordance with the requirements of MIT’s Committee on the Use of Humans as Experimental Subjects (COUHES) and were paid.

#### Study 1

Thirty-two individuals with a clinical ASD diagnosis participated. Four participants were not included in the analyses due to motion-related scanner artifacts, leaving 28 participants. All participants were native English speakers with normal hearing and vision (*M*_age_=26.5, *SD*=6.5, range: 18-45; 7 females; 5 left-handed). All participants were administered the Autism Diagnostic Observation Schedule (ADOS; Lord et al., 1999) and met the criteria for a clinical diagnosis: *M*=9.6, *SD*=2.3 (these summary statistic exclude 2 participants whose ADOS scores were lost). All participants were also administered the Autism Spectrum Quotient questionnaire (ASQ; Baron-Cohen et al., 2001): *M*=31.7, *SD*=8.4. Finally, nonverbal IQ was measured with the Kaufman Brief Intelligence Test (KBIT; Kaufman, 1990): *M*=113.8, *SD*=11.9.

Twenty-eight native English speakers without a clinical ASD diagnosis or any other neurodevelopmental disorder were pairwise-matched to ASD participants on age (*M*=25.8, *SD*=5.3, range: 20-45), sex, handedness, nonverbal IQ (*M*=118.1, *SD*=9.8), fMRI acquisition-sequence parameters, experimental parameters (the version of the localizer used for each network, as detailed below), and the amount of motion in the scanner during each task (**Table 1**; see Jenkinson (1999) for details on motion measurements). The ASQ questionnaire confirmed that none of the participants exhibited clinically significant levels of autistic traits: *M*=18.0, *SD*=6.6.

**Table 1.** Matching the ASD and NT participants (N=28 in each group) for Study 1.

|  | Group |  | *t*-test |
| --- | --- | --- | --- |
|  | ASDs | NTs |  |
| Sex (male:female) | 21:7 | 21:7 |  |
| Handedness (right:left) | 23:5 | 23:5 |  |
| Age: Mean (SD) | 26.5 (6.5) | 25.8 (5.5) | *t*(54) = 0.46 |
| ADOS: Mean (SD) | 9.6 (2.3) | NA |  |
| ASQ: Mean (SD) | 31.7 (8.4) | 18.0 (6.6) | ***t*(51) = 6.62\*** |
| KBIT: Mean (SD) | 113.8 (11.9) | 118.1 (9.8) | *t*(54) = 1.43 |
| TVM - Language Localizer task: Mean (SD) | 0.14 (0.06) | 0.12 (0.03) | *t*(54) = 1.63 |
| TVM - ToM Localizer task: Mean (SD) | 0.14 (0.05) | 0.13 (0.04) | *t*(43) = 0.48 |
| TVM - MD Localizer task: Mean (SD) | 0.13 (0.05) | 0.12 (0.03) | *t*(36) = 1.05 |

Notes: TVM – total vector motion (in mm). Results of *t*-tests significant at $p < 0.05$ are bolded and marked by an asterisk.

#### (Exploratory) Study 2

We searched the Fedorenko lab’s database for native English speakers with normal hearing and vision, no ASD diagnosis (or other neurodevelopmental/psychiatric disorders), who had completed the ASQ and KBIT. These criteria yielded 189 individuals (*M*_age_=26.2, *SD*=6.6, range: 19-61; 132 females; 14 left-handed; *M*_ASQ_=17.3, *SD*=7.2, range: 3-42; *M*_KBIT_=119.3, *SD*=11.9, range: 75-132). Eight participants (4%) had clinically significant levels of autistic traits (ASQ score ≥ 32).

### 2.2. Design, materials, and procedure

#### Study 1

All participants completed a language localizer (Fedorenko et al., 2010). A subset of participants additionally completed a Theory of Mind localizer (Saxe & Kanwisher, 2003) (n=18 in each group), and/or a Multiple Demand system localizer (e.g., Blank et al., 2014) (n=19 in each group). (Not everyone completed all three localizers because this dataset was pooled from across two projects differing in their goals.) Some participants completed additional tasks for unrelated studies. The scanning session lasted approximately 2 hours.

##### Language localizer

Participants read sentences nonword-sequences in a blocked design. The Sentences>Nonwords contrast targets brain regions that selectively support highlevel linguistic processing, including lexical-level and combinatorial—syntactic and semantic—processes (Fedorenko et al., 2010; 2012; 2018). Two slightly different versions of the localizer were used (for the reason noted above), which have been previously established to elicit similar activations (Fedorenko et al., 2010; 2014). In one version (*SNloc_ips189*), each trial started with 300 ms pre-trial fixation, followed by a 12-word-long sentence/nonword-sequence presented one word/nonword at a time for 350ms each, followed by a probe word/nonword presented in blue for 1000ms. Participants pressed one of two buttons to indicate whether the probe appeared in the preceding stimulus. Each trial ended with 500ms of fixation. In the other version (*SNloc_ips179*), each trial started with 100ms pre-trial fixation, followed by a 12-word-long sentence/nonword-sequence presented at the rate of 450ms per word/nonword, followed by a line drawing of a finger pressing a button presented for 400ms. Participants pressed a button whenever they saw this drawing. Each trial ended with 100ms of fixation. In both versions, each block consisted of 3 trials and lasted 18s. Each run consisted of 16 experimental blocks (8 per condition) and 5 fixation blocks (18s or 14s each for the two versions), for a total duration of 378s or 358s. Each participant performed two runs, with condition order counterbalanced across runs.

##### ToM localizer

Participants read short stories. In the critical, False Belief, condition, each story described a protagonist who held a false belief. In the control, False Photograph, condition, each story described an object (e.g., a photograph or a painting) depicting some state of the world that was no longer true. The False Belief>False Photo contrast targets brain regions that support Theory of Mind reasoning (Dodell-Feder et al., 2011; Saxe & Kanwisher, 2003). The stories were presented one at a time for 10s, centered on the screen, and followed by a True/False question presented for 4s. Participants pressed one of two buttons to indicate their response. Each run consisted of 10 trials (5 per condition), and 11 fixation blocks, for a total duration of 272s. Each participant performed two runs, with condition order counterbalanced across runs.

##### MD localizer task

Participants kept track of four (easy condition) or eight (hard condition) spatial locations in a 3×4 grid (Fedorenko et al., 2011). The Hard>Easy contrast targets brain regions that support domain-general executive processes, like working memory and cognitive control (Duncan, 2010, 2013; Fedorenko et al., 2013). The locations flashed up one or two at a time (for the easy and hard conditions, respectively), followed by the presentation of two sets of locations. Participants pressed one of two buttons to indicate which set of locations they just saw. Each trial lasted 8 s (see Fedorenko et al., 2011, for details). Each block consisted of 4 trials and lasted 32s. Each run consisted of 12 experimental blocks (6 per condition) and 4 fixation blocks (16s each), for a total duration of 448s. Each participant performed two runs, with condition order counterbalanced across runs.

#### Study 2

Participants completed one of six versions of the language localizer task (**Table 2**), with 165 of the 189 participants (87%) completing the versions used in Study 1. Detailed information on the procedure and timing of the different versions can be found in Mahowald and Fedorenko (2016).

**Table 2.** Information on which subsets of participants in the sample of 189 NT participants performed which version of the language localizer. Information on the procedure and timing details for the SNloc_ips179 and SNloc_ips189 is provided in the Methods section. Information on the procedure and timing details for the other versions of the language localizer can be found in Mahowald & Fedorenko (2016), Table 2.

| Number of participants | Language localizer version | Conditions | Materials | Trials per block | Blocks per run / per condition per run |
| --- | --- | --- | --- | --- | --- |
| n = 153 | SNloc_ips179 | Sentences, Nonwords | 12-word-/nonword-long sequences | 3 | 16/8 |
| n = 12 | SNloc_ips189 | Sentence, Nonwords | 12-word-/nonword-long sequences | 3 | 16/8 |
| n = 13 | SWNloc_ips168 | Sentences, Wordlists, Nonwords | 8-word-/nonword-long sequences | 5 | 12/4 |
| n = 6 | SWNloc_ips198 | Sentences, Wordlists, Nonwords | 12-word-/nonword-long sequences | 3 | 18/6 |
| n = 1 | SWJN_v1_ips252 | Sentences, Wordlists, Jaberwocky, Nonwords | 12-word-/nonword-long sequences | 5 | 16/4 |
| n = 3 | SWJN_v2_ips232 | Sentences, Wordlists, Jaberwocky, Nonwords | 8-word-/nonword-long sequences | 5 | 16/4 |
| n = 1 | SNloc_ips232 | Sentences, Nonwords | 8-word-/nonword-long sequences | 5 | 16/8 |

### 2.3. fMRI data acquisition and preprocessing

Structural and functional data were collected on the whole-body 3 Tesla Siemens Trio scanner with a 32-channel head coil at the Athinoula A. Martinos Imaging Center at the McGovern Institute for Brain Research at MIT. T1-weighted structural images were collected in 128 axial slices with 1mm isotropic voxels (TR=2,530ms, TE=3.48ms). Functional, blood oxygenation level dependent (BOLD) data were acquired using an EPI sequence (with a 90° flip angle and using GRAPPA with an acceleration factor of 2), with the following acquisition parameters: thirty-one 4mm thick near-axial slices, acquired in an interleaved order with a 10% distance factor; 2.1mm × 2.1mm in-plane resolution; field of view 200mm in the phase-encoding anterior to posterior direction; matrix size 96mm × 96mm; TR 2,000ms; TE 30ms; 16 nonlinear iterations for spatial normalization with 7×9×7 basis functions. Prospective acquisition correction (Thesen et al., 2000) was used to adjust the positions of the gradients based on the participant’s motion one TR back. The first 10s of each run were excluded to allow for steady-state magnetization.

MRI data were analyzed using SPM5 and custom MATLAB scripts. Each participant’s data were motion corrected and normalized into a common brain space (the Montreal Neurological Institute, MNI, brain template) and resampled into 2mm isotropic voxels. The data were then smoothed with a 4mm Gaussian filter and high-pass filtered (at 200s). The effects were estimated using a General Linear Model (GLM) in which each experimental condition was modeled with a boxcar function (representing entire blocks/events) convolved with the canonical hemodynamic response function. The model also included first-order temporal derivatives of these effects, as well as nuisance regressors representing experimental runs and offline-estimated motion parameters.

### 2.4. Definition of group-constrained, subject-specific fROIs

For each participant, functional ROIs (fROIs) were defined using the Group-constrained Subject-Specific (GSS) approach (Fedorenko et al., 2010). In this approach, a set of parcels (brain areas within which most individuals in prior studies showed activity for the relevant localizer contrast) is combined with each individual participant’s activation map for the same contrast. To define the language fROIs in the LH, we used parcels—derived from a probabilistic activation overlap map for the Sentences>Nonwords contrast for 220 participants—falling within inferior frontal gyrus (LIFG) and its orbital part (LIFGorb), middle frontal gyrus (LMFG), anterior and posterior temporal (LAntTemp and LPostTemp), and angular gyrus (LAngG). Further, we defined the RH homologous fROIs using LH parcels mirror-projected onto the RH. The mirrored versions of the parcels are likely to encompass the RH homolog of the LH language network, despite possible hemispheric asymmetries in their precise locations.

For the ToM network, we focused on the RTPJ, which has been shown to be most selective for mental state attribution (Saxe & Powell, 2006) and its LH homolog. To define these fROIs, we used the parcels derived from a group-level representation for the False Belief>False Photo contrast in an independent group of 462 participants (Dufour et al., 2013).

To define the MD fROIs, following Fedorenko et al. (2013), we used eighteen anatomical parcels across the two hemispheres (Tzourio-Mazoyer et al., 2002): opercular IFG (LIFGop & RIFGop), MFG (LMFG & RMFG), orbital MFG (LMFGorb & RMFGorb), insular cortex (LInsula & RInsula), precentral gyrus (LPrecG & RPrecG), supplementary and presupplementary motor areas (LSMA & RSMA), inferior parietal cortex (LParInf & RParInf), superior parietal cortex (LParSup & RParSup), and anterior cingulate cortex (LACC & RACC).

These parcels were used to extract functional *region volumes* and *effect sizes* in each network in each individual. To compute *region volumes*, we counted the number of voxels showing a significant effect (at the *p*<0.001 whole-brain uncorrected threshold) for the relevant localizer contrast within each relevant parcel. For example, for the language network, we counted the number of significant Sentences>Nonwords voxels within each of six LH and each of six RH language parcels. For the language and MD networks, these values were then summed across the regions within each hemisphere, to derive a single value per hemisphere per network per participant. The motivation for examining the networks holistically (cf. region by region) is that much evidence suggests that the regions within each network form a strongly functionally integrated system (Blank et al., 2014; Mahowald & Fedorenko, 2016; Fedorenko & Blank, in press).

To compute *effect sizes*, we first defined subject-specific fROIs by selecting the top 10% of most localizer-responsive voxels based on the *t*-values for the relevant contrast and then extracted the responses of these fROIs to the relevant localizer contrast (in percent BOLD signal change). To ensure independence between the data used to define the fROIs vs. to extract effect-size measures (Kriegeskorte et al., 2009), we used an across-runs cross-validation procedure (Nieto-Castañon & Fedorenko, 2012). In particular, the first run was used to define the fROIs, and the second run to estimate the responses, the second run was used to define the fROIs, and the first run to estimate the responses, and finally, the estimates were averaged across the two left-out runs to derive a single value per participant per fROI. For the language and MD networks, these values were then averaged across the regions within each hemisphere, to derive a single value per hemisphere per network per participant.

### 2.5. Computing lateralization

The critical measure for each network was the degree of *lateralization*. Following prior work (Binder et al., 1997; Seghier et al., 2008), we used volume-based lateralization (cf. activation-strength-based lateralization; see Mahowald & Fedorenko, 2016, for evidence that the two are correlated). For each network, the number of activated voxels in the RH was subtracted from the number of activated voxels in the LH, and the resulting value was divided by the number of activated voxels across hemispheres. The obtained values could therefore range from 1 (exclusively LH activation) to −1 (exclusively RH activation), with 0 corresponding to bilateral activations. We have previously established that this measure is highly stable within individuals over time (Mahowald & Fedorenko, 2016).

### 2.6. Analyses

#### Study 1

We compared ASD individuals to pairwise-matched NT controls. To test for group differences in language lateralization, we used GLMs with *Group* (ASD vs. NT) as a predictor of lateralization in the language network identified using the Sentences>Nonwords contrast of the language localizer task. We also examined data fit under the null and alternative hypotheses by estimating Bayes factors (BF10) using Bayesian linear regressions (Wagenmakers et al., 2011). BF10 statistics—which reveal how many times the observed data are more likely under the alternative than the null hypothesis—were calculated using the JASP software package. We adopted the following interpretations of BF10 (Lee and Wagenmakers, 2013): BF10<1 provides evidence for the null hypothesis, 1<BF10<3 is considered anecdotal evidence for the alternative hypothesis, 3<BF10<10—moderate evidence, and 10<BF10—strong evidence.

To determine whether presumed differences in language lateralization result from differences in the LH, the RH, or both, we submitted LH and RH *region volumes*, as well as *effect sizes* (in a separate analysis), as dependent variables to GLMs and Bayesian linear regressions with *Group* (ASD vs. NT) as a predictor. Effect sizes are generally highly correlated with volumes but show greater stability at the individual level given that they do not depend on the statistical-significance threshold (Mahowald & Fedorenko, 2016). In an additional analysis, to assess the robustness of the language lateralization differences across different linguistic materials, we further examined the responses in the language fROIs to the processing of the False Photo stories from the ToM localizer (relative to fixation).

To test whether lateralization reduction is restricted to the language network, we submitted LH and RH *region volumes* and *effect sizes* as dependent variables to GLMs and Bayesian linear regressions with *Group* (ASD vs. NT) as a predictor of lateralization in the ToM and MD networks. We also computed bivariate and partial (controlling for age and nonverbal IQ) Pearson (here and elsewhere) correlations between language and MD lateralization measures (networks where lateralization reduction was observed in individuals with ASD) across participants, to test whether these effects are driven by the same underlying factor.

Finally, to test whether language lateralization in our sample of individuals with ASD is related to autism severity, we computed bivariate and partial (controlling for age and non-verbal IQ) correlations between language lateralization measures and ASQ and ADOS scores.

#### (Exploratory) Study 2

To test whether language lateralization is related to autistic trait load in NT individuals, we computed bivariate and partial (controlling for age and non-verbal IQ) correlations between language lateralization measures and ASQ scores.

## 3. Results

### 3.1. Language lateralization is reduced in ASD individuals

Language activations were left-lateralized in both groups (Sentences>Nonwords: *M*_*ASD*_=0.36, *SD*=0.06; *M*_*NT*_=0.61, *SD*=0.04; False Photo>Fixation: *M*_*ASD*_=0.16, *SD*=0.48; *M*_*NT*_=0.54, *SD*=0.18). Critically, however, the degree of lateralization was significantly lower in ASD individuals (Sentences>Nonwords: *t*(54)=2.90, *p*=0.005, BF10=7.71; False Photo>Fixation: *t*(34)=3.10, *p*=0.004, BF10=10.46; **Figure 1a**). This difference was primarily driven by differences in the RH activity (**Figures 1b-c**). In particular, for the Sentences>Nonwords contrast, LH region volumes and effect sizes were similar between the groups (volumes: *t*(54)=0.56, *p*=0.58, BF10=0.31; effect sizes: *t*(54)=1.19, *p*=0.24, BF10=0.49), but in the RH, ASD individuals exhibited larger volumes (*t*(54)=-2.44, *p*=0.018, BF10=3.10) and effect sizes (*t*(54)=-2.94, *p*=0.005, BF10=8.47) than controls. Similarly, for the False Photo>Fixation contrast, LH volumes were similar between the two groups (*t*(34)=1.56, *p*=0.13, BF10=0.82), but in the RH, ASD individuals exhibited somewhat larger volumes (*t*(34)=-2.22, *p*=0.03, BF10=2.07), although no reliable group differences obtained for the effect sizes (LH: *t*(34)=1.83, *p*=0.08; BF10=1.15; RH: *t*(34)=-0.91, *p*=0.37, BF10=0.44).

**Figure 1.**
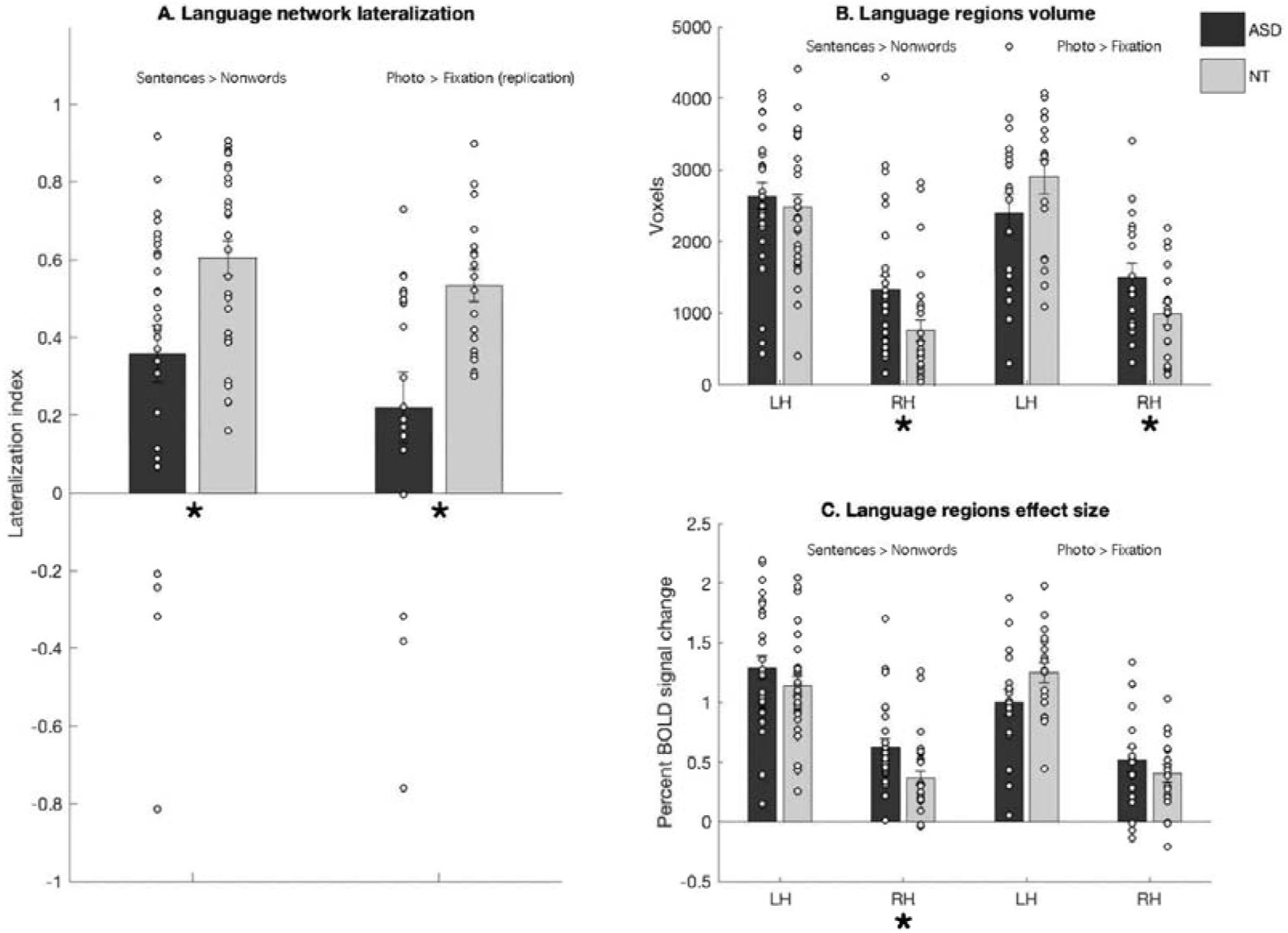
**A.** Lateralization measures for the language network in individuals with ASD (n=28, darker bars) vs. NT controls (n=28, lighter bars). **B**. Region volumes of the language LH and RH networks. **C.** Effect sizes of the language LH and RH networks. Significant group differences are marked by asterisks.

### 3.2. Reduced language lateralization in ASD does not stem from generalized reduction in lateralization across the brain

ToM activations were right-lateralized in both groups (*M*_*ASD*_=-0.15, *SD*=0.37; *M*_*NT*_=-0.12, *SD*=0.37). Replicating Dufour et al. (2013), we found no group difference in lateralization (*t*(34)=0.21, *p*=0.84, BF10=0.33; **Figure 2a**). Further, also in line with Dufour et al. (2013), the groups did not reliably differ in the volumes or effect sizes in either the LH or RH (LH volumes: *t*(34)=0.74, *p*=0.47, BF10=0.40; LH effect sizes: *t*(34)=0.38, *p*=0.71, BF10=0.34; RH volumes: *t*(34)=0.03, *p*=0.98, BF10=0.32; RH effect sizes: *t*(34)=0.08, *p*=0.94, BF10=0.32; **Figure 2b-c**).

**Figure 2.**
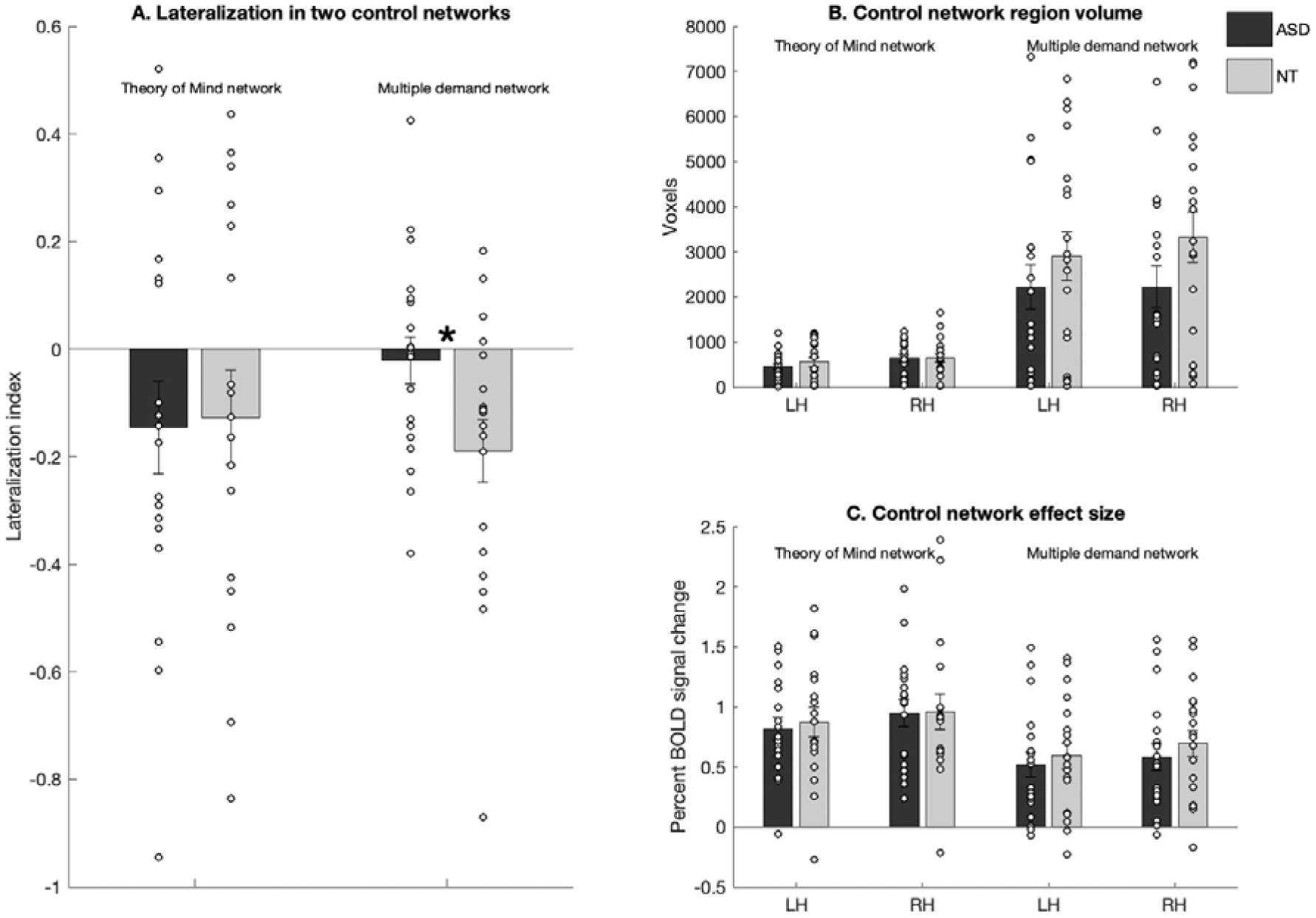
**A.** Lateralization measures for the ToM and MD networks in individuals with ASD (n=28, darker bars) vs. NT controls (n=28, lighter bars). **B**. Region volumes of the ToM and MD LH and RH networks. **C.** Effect sizes of the ToM and MD LH and RH networks. Significant group differences are marked by asterisks.

MD activations did not show a laterality bias in ASD participants (*M*_*ASD*_=-0.02, *SD*=0.19), and they were right-lateralized in the control group (*M*_*NT*_=-0.19, *SD*=0.25). The degree of lateralization was lower in ASD individuals (*t*(36)=2.30, *p*=0.03; **Figure 2a**), although according to a Bayes factor (BF10=2.37), this evidence was weak. Further, no reliable group differences obtained for the volumes or effect sizes in either the LH or RH (LH region volumes: *t*(36)=0.92, *p*=0.36, BF10=0.44; LH effect sizes: *t*(36)=0.51, *p*=0.62, BF10=0.35; RH region volumes: *t*(36)=1.52, *p*=0.14, BF10=0.77; RH effect sizes: *t*(36)=0.71, *p*=0.481; BF10=0.39; **Figure 2b-c**).

Given that ASD individuals showed reduced lateralization in the language network and, to some extent, in the MD network, we asked whether the degree of this reduction was correlated between the two networks, which would suggest a shared underlying mechanism. The lateralization measures showed no reliable correlation (*r*(17)=0.09, *p*=0.71; partial *r*(15)=0.11, *p*=0.67), suggesting that these effects are independent.

### 3.3. Reduced language lateralization relates to autistic trait load

In 189 NT participants, ASQ scores correlated significantly with the degree of language lateralization, with higher autistic trait load associated with less lateralized responses (*r*(187)=-0.15, *p*=0.05, partial *r*(185)=-0.15, *p*=0.04; **Figure 3b**). A further exploratory examination of the correlations between the lateralization measure and ASQ subscale scores revealed that the observed relationship is primarily driven by the communicative abilities subscale (*r*(187)=-.18, *p*=.05, partial *r*(185) =-.20, *p*=.04; p-values are adjusted using a Bonferroni correction to account for multiple comparisons (n=5 subscales)). Scores on the other subscales—tapping social skills, imagination, attention switching, or attention to detail—did not reliably correlate with language lateralization (all *rs*(187)<|0.11|, all *p*s>0.14; all partial *rs*(187)<|0.11|, all *p*s>0.15).

**Figure 3.**
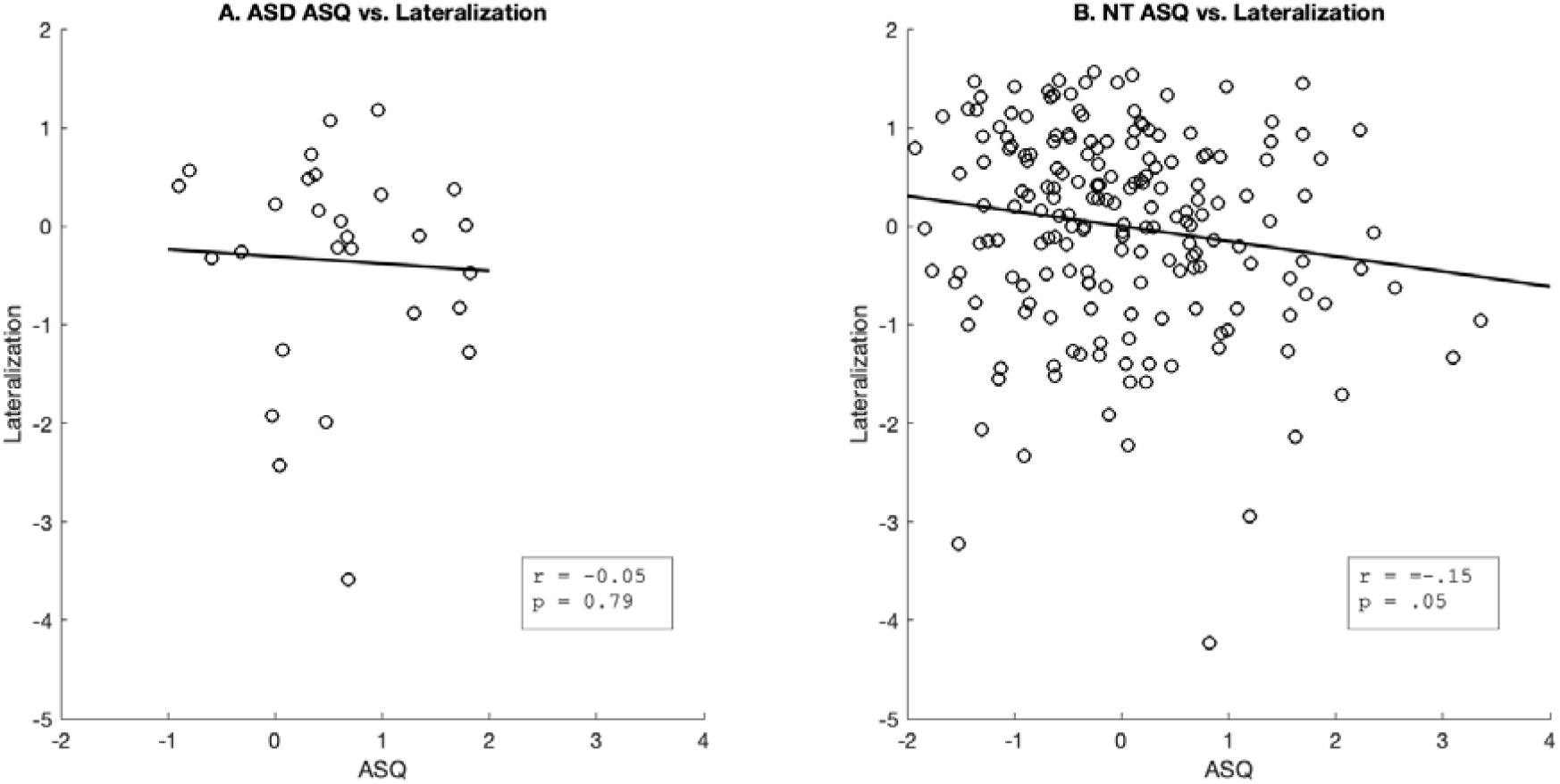
**Reduced language lateralization and the presence of autistic traits. A:** A correlation between the language lateralization measure and autism severity, as measured by ASQ scores, in individuals with ASD (n=28). **B**: A correlation between the language lateralization measure and autistic trait load, as measured by the ASQ scores, in NTs (n=189). The datapoints are standardized residual values, controlling for age and nonverbal IQ.

We also examined relations between language lateralization and ASQ or ADOS scores in the ASD participants keeping in mind that correlations in small samples should be interpreted with caution (Schönbrodt & Perugini, 2013). Neither ASQ scores nor ADOS scores correlated with the degree of lateralization (ASQ: *r*(26)=-.43, *p*=.02; partial *r*(24)=-.34; *p=*.09; **Figure 3a;** ADOS: *r*(24)=-.16, *p*=.44; partial *r*(22)=-.04; *p*=.85), possibly due to insufficient variability in the ASQ and ADOS scores in this population (Hedge et al., 2018).

## 4. Discussion

Perhaps the most consistent finding from cognitive neuroscience of autism is reduced lateralization of neural activity during language tasks (Lindell & Hudry, 2013; Herringshaw et al., 2016). However, as discussed in the Introduction, most prior studies have relied on comparisons of group-level maps, complicating interpretation, and left the nature and scope of this effect ambiguous. Here, using a robust individual-subjects functional localization approach (Fedorenko et al., 2010), we found reduced lateralization during language comprehension—across two paradigms—in individual participants. Furthermore, we established that this reduced lateralization (i) results from increased RH activity (cf. decreased LH activity or both); (ii) occurs within the language-selective network (given that the language localizer effectively isolates this network from nearby functionally distinct networks; Braga et al., 2019; Fedorenko & Blank, in press); (iii) is not due to a global lateralization reduction (given that two other lateralized networks did not show a similar reduction). Finally, in a more exploratory investigation of neurotypical individuals, we found some evidence for reduced language lateralization in individuals with high autistic trait load. Below, we discuss several issues that our results bear on, and highlight some open questions and challenges for future work.

### 4.1. Speech/language lateralization in typical development

Long before they understand or utter their first words, typically developing infants have been argued to show lateralized responses to speech (Dahaene-Lambertz et al., 2002; Pena et al., 2003; Minagawa-Kawai et al., 2011), although the selectivity of this bias for speech relative to non-speech sounds has been questioned (Christia et al., 2014). This early, possibly innate, LH bias for speech perception presumably drives the lateralization of the later-emerging system that supports language comprehension, which is in place certainly by age ∼4-5 years—as early as we have been able to measure with fMRI (Berl et al., 2014; Friederici et al., 2011; Holland et al., 2007) and likely, earlier, given what we know about language development from behavioral work (Bloom, 1968). However, there is an unresolved paradox surrounding developmental changes in speech and language lateralization. In particular, unlike high-level comprehension processes—the focus of the current investigation—which are strongly left-lateralized in adults, speech perception appears to be bilateral in adulthood (Peele, 2012; Poeppel, 2014; Norman-Haignere et al., 2015). So, the alleged innate/early LH bias for speech perception apparently disappears as the brain matures and/or gets more linguistic experience. High-level comprehension, on the other hand, shows the opposite pattern: activations tend to be more bilateral at younger ages, becoming more strongly left-lateralized toward adulthood (Berl et al., 2014; Szaflarski et al., 2006; McNealy et al., 2011). So, the emergence of speech—and later, language—lateralization in typical development remains to be characterized more precisely and understood at a functional level (in terms of the behavioral/cognitive advantages it confers) before we can meaningfully probe deviations in its time-course and other properties in developmental disorders.

### 4.2. Increased RH activity during language processing in ASD

We found stronger RH responses during language processing in ASD individuals compared to controls, leading to more bilateral responses. Is the LH bias for speech absent/reduced in autism and other developmental disorders at birth, or is the LH bias ubiquitous at birth, with the RH playing a gradually stronger role in individuals who experience speech/language difficulties (Bishop, 2013)?

Some have proposed that increased RH activity stems from an aberrant brain development trajectory in autism (Courchesne et al., 2001; Redcay & Courchesne, 2005). In particular, whereas typically developing brains grow at a relatively constant rate (Toga & Thompson, 2003), brains of ASD individuals exhibit initial rapid growth, followed by a premature arrest of growth (Courchesne, 2004; Pardo & Eberhart, 2007). Because the RH matures earlier than the LH (Chiron et al., 1997; Geschwind et al., 2002), this difference in the growth rate at different brain development stages may lead to an over-developed RH and under-developed LH in ASD (Courchesne, 2004). Furthermore, genes related to language and cognitive development, like *FOXP2* (Fisher & Scharff, 2009; MacDermot et al., 2005) or *CNTNAP2* (Alarcon et al., 2008; Scott-Van Zeeland et al., 2010), are differentially expressed in the left vs. right embryonic perisylvian cortex (Sun et al., 2005). Dysregulation of these genes in autism (Muhle et al., 2004; cf. Newbury et al., 2002) could potentially lead to increased RH engagement during language processing. Others have advanced an experiential account whereby speech/language difficulties in ASD individuals result in overtaxing the specialized LH mechanisms (cf. the finding noted above of early bilateral language responses in NT children) and the consequent recruitment of less specialized homologous RH areas (Mason et al., 2008; Tesink et al., 2009).

Regardless of the origin of increased RH activity in autism, we can ask whether it leads to better language outcomes, or is maladaptive. There are both theoretical and empirical reasons in support of the latter possibility. On the theoretical side, RH language regions have been argued to be less functionally specialized for language processing (Gotts et al., 2013; Lindell, 2006), instead supporting visual semantic processing and/or visual imagery (Joseph, 1988; Roland & Friberg, 1985). Indeed, some individuals with ASD describe themselves as “visual thinkers”, who translate linguistic representations into mental images to achieve comprehension (Grandin, 2006). Some have further proposed that RH language regions are less engaged in linguistic prediction (Federmeier & Kutas, 1999; Federmeier et al., 2008). And empirically, in aphasia research, the recruitment of RH language regions (cf. the intact LH language regions) following stroke has been argued to be maladaptive (Barwood et al., 2011; Hamilton et al., 2010; Turkeltaub et al., 2012).

Our data also suggestively point to the maladaptive nature of the RH engagement: NT individuals with more bilateral language responses reported greater communication difficulties, and a similar trend was present in ASD individuals, although the sample is too small to draw meaningful conclusions. So, greater RH activity does not appear to alleviate language/communicative difficulties.

### 4.3. Evidence against global lateralization reduction in ASD

Some have argued that reduced lateralization in autism extends beyond language processing (Dawson, 1983; Fein et al., 1984; Cardinale et al., 2013; Postema et al., 2019). To test this idea, in addition to the language network, we examined activity in two other networks that are right-lateralized in NT individuals: the ToM mentalizing network (Saxe & Kanwisher, 2003) and the domain-general Multiple Demand (MD) executive control network (Duncan, 2010). We found no evidence for decreased lateralization of the ToM activity, and only weak evidence for decreased lateralization of the MD activity in ASD individuals (see also Dufour et al., 2013; Gilbert et al., 2006). These results argue against a generally more functionally symmetrical brain in ASD individuals: reduced lateralization appears to be most pronounced—and perhaps restricted to—the language network (see also Nielsen et al., 2014, for similar conclusions drawn from resting state fMRI).

### 4.4. Reduced language lateralization as a feature of the broader autism phenotype

Autism-like traits are present to some degree in many individuals who do not have a clinical ASD diagnosis. Subclinical presentation of such traits, known as the broader autism phenotype (BAP), is thought to index genetic liability for this neurodevelopmental disorder (Landry & Chouinard, 2016; Piven et al., 1997). In our exploratory Study 2, we observed reduced language lateralization in NT individuals with a higher autistic trait load. These results demonstrate that reduced language lateralization may extend to NT individuals with increased risk for autism, in line with a continuum model of underlying genetic risk (Wing, 1988; Geschwind, 2011; Gaugler et al., 2014; Robinson et al., 2016).

### 4.5. Limitations and Future Directions

Functional brain-imaging (and many behavioral) investigations of ASD are characterized by small samples, which is problematic given the well-documented heterogeneity of this population. Furthermore, most studies, including ours, only include high-functioning individuals with ASD (cf. Tager-Flusberg & Kasari, 2013). Yet lower-functioning ASD individuals, some of whom never acquire functional linguistic skills (Maljaars et al., 2012), may hold critical clues as to the nature and neural basis of the linguistic/communicative impairment in ASD.

Another challenge—relevant to probing the *functional importance* of reduced language lateralization—is the lack of robust and validated behavioral measures of linguistic processing that are not confounded by executive demands, like commonly used vocabulary and grammar assessments that strongly correlate with measures of non-verbal IQ (Beck & Black, 1986; Hodapp & Gerken, 1999). Efforts to develop robust language-selective measures will be critical in understanding how more bilateral language processing affects behavioral outcomes.

Finally, evidence for reduced language lateralization has been previously provided for diverse developmental disorders, including those that affect linguistic functions, like dyslexia and specific language impairment, but also those that do not typically affect language/communication, like schizophrenia, attention-deficit hyperactivity disorder, and epilepsy (De Guibert et al., 2011; Hale et al., 2005; Oertel-Knöchel & Linden, 2011; Wehner et al., 2007; Yuan et al., 2006). Whether patterns of atypical lateralization for language differ across these disorders, and whether they result from the same underlying mechanisms remains to be determined.

## Acknowledgements

We would like to acknowledge the Athinoula A. Martinos Imaging Center at McGovern Institute for Brain Research at MIT, and the support team (Steve Shannon and Atsushi Takahashi). This work was supported by a grant from the Simons Foundation to the Simons Center for the Social Brain at MIT to EF and NK. EF was additionally supported by NIH awards R00-HD057522 and R01-DC016607. We also thank Matt Siegelman for help with the figures, and Rebecca Saxe for helpful discussions.

## References

Alarcón, M., Abrahams, B. S., Stone, J. L., Duvall, J. A., Perederiy, J. V., Bomar, J. M., … & Nelson, S. F. (2008). Linkage, association, and gene-expression analyses identify CNTNAP2 as an autism-susceptibility gene. The American Journal of Human Genetics, 82(1), 150–159.

Amunts, K., Schleicher, A., Bürgel, U., Mohlberg, H., Uylings, H. B. M., & Zilles, K. (1999). Broca’s region revisited: Cytoarchitecture and intersubject variability. Journal of Comparative Neurology, 412(2), 319–341.

Anderson, J. S., Lange, N., Froehlich, A., Dubray, M. B., Druzgal, T. J., Froimowitz, M. P., … Lainhart, J. E. (2010). Decreased left posterior insular activity during auditory language in autism. American Journal of Neuroradiology, 31(1), 131–139.

Baird, G., Charman, T., Baron-Cohen, S., Cox, a, Swettenham, J., Wheelwright, S., & Drew, A. (2000). A screening instrument for autism at 18 months of age: a 6-year follow-up study. Journal of the American Academy of Child and Adolescent Psychiatry, 39(6), 694–702.

Baron-Cohen, S., Lombardo, M. V., Auyeung, B., Ashwin, E., Chakrabarti, B., & Knickmeyer, R. (2011). Why are autism spectrum conditions more prevalent in males?. PLoS biology, 9(6), e1001081.

Barwood C.H., Murdoch B.E., Whelan B.M., Lloyd D., Riek S., Jd O.S., Coulthard A., Wong A. (2011). Improved language performance subsequent to low-frequency rTMS in patients with chronic non-fluent aphasia post-stroke. European Journal of Neurology, 18(7), 935–943.

Beck, F. W., & Black, F. L. (1986). Comparison of PPVT—R and WISC—R in a Mild/Moderate Handicapped Sample. Perceptual and motor skills, 62(3), 891–894.

Berl, M. M., Mayo, J., Parks, E. N., Rosenberger, L. R., VanMeter, J., Ratner, N. B., … & Gaillard, W. D. (2014). Regional differences in the developmental trajectory of lateralization of the language network. Human Brain Mapping, 35(1), 270–284.

Binder, J. R., Frost, J., Hammeke, T., Cox, R.W., Rao, S.M., & Prieto, T. (1997). Human brain language areas identified by functional magnetic resonance imaging. The Journal of Neuroscience, 17(1), 353–362.

Bishop, D.V. (2013). Cerebral asymmetry and language development: cause, correlate, or consequence?. Science, 340(6138), 1230531.

Blank, I., Balewski, Z., Mahowald, K., & Fedorenko, E. (2016). Syntactic processing is distributed across the language system. NeuroImage, 127, 307–323.

Blank, I., Kanwisher, N., Fedorenko, E., (2014). A functional dissociation between language and multiple-demand systems revealed in patterns of BOLD signal fluctuations. Journal of Neurophysiology, 112, 1105–1118.

Bloom, L. M. (1968). Language development: Form and function in emerging grammars.

Boddaert, N., Belin, P., Chabane, N., Poline, J. B., Barthélémy, C., Mouren-Simeoni, M. C., … Zilbovicius, M. (2003). Perception of complex sounds: Abnormal pattern of cortical activation in autism. American Journal of Psychiatry, 160(11), 2057–2060.

Bradshaw, A.R., Thompson, P.A., Wilson, A.C., Bishop, D.V., & Woodhead, Z.V. (2017). Measuring language lateralisation with different language tasks: a systematic review. PeerJ, 5, e3929.

Braga, R.M., Van Dijk, K.R., Polimeni, J.R., Eldaief, M.C., & Buckner, R.L. (2019). Parallel distributed networks resolved at high resolution reveal close juxtaposition of distinct regions. Journal of neurophysiology, 121(4), 1513–1534.

Cardinale, R.C., Shih, P., Fishman, I., Ford, L.M., & Müller, R.-A. (2013). Pervasive Rightward Asymmetry Shifts of Functional Networks in Autism Spectrum Disorder. JAMA Psychiatry, 70(9), 975.

Chiron, C., Jambaque, I., Nabbout, R., Lounes, R., Syrota, A., & Dulac, O. (1997). The right brain hemisphere is dominant in human infants. Brain, 120(6), 1057–1065.

Cristia, A., Minagawa, Y., & Dupoux, E. (2014). Responses to vocalizations and auditory controls in the human newborn brain. PLoS One, 9(12).

Courchesne, E. (2004). Brain development in autism: early overgrowth followed by premature arrest of growth. Mental retardation and developmental disabilities research reviews, 10(2), 106–111.

Courchesne, E., Karns, C.M., Davis, H.R., Ziccardi, R., Carper, R.A., Tigue, Z.D., … Courchesne, R.Y. (2001). Unusual brain growth patterns in early life in patients with autistic disorder: An MRI study. Neurology, 57(2), 245–254.

Dawson, G. (1983). Lateralized brain dysfunction in autism: Evidence from the Halstead-Reitan neuropsychological battery. Journal of Autism and Developmental Disorders, 13(3), 269–286.

Deen, B., & Pelphrey, K. (2012). Perspective: Brain scans need a rethink. Nature, 491(7422), S20–S20.

De Guibert, C., Maumet, C., Jannin, P., Ferré, J. C., Tréguier, C., Barillot, C., … Biraben, A. (2011). Abnormal functional lateralization and activity of language brain areas in typical specific language impairment (developmental dysphasia). Brain, 134(10), 3044–3058.

Dehaene-Lambertz, G., Hertz-Pannier, L., & Dubois, J. (2006). Nature and nurture in language acquisition: anatomical and functional brain-imaging studies in infants. Trends in Neurosciences.

Dufour, N., Redcay, E., Young, L., Mavros, P. L., Moran, J. M., Triantafyllou, C., … Saxe, R. (2013). Similar Brain Activation during False Belief Tasks in a Large Sample of Adults with and without Autism. PLoS ONE, 8(9).

Duncan, J. (2010). The multiple-demand (MD) system of the primate brain: mental programs for intelligent behaviour. Trends in Cognitive Science, 14, 172–179.

Duncan, J. (2013). The structure of cognition: attentional episodes in mind and brain. Neuron, 80, 35–50.

Eyler, L.T., Pierce, K., & Courchesne, E. (2012). A failure of left temporal cortex to specialize for language is an early emerging and fundamental property of autism. Brain, 135(3), 949–960.

Federmeier, K.D., & Kutas, M. (1999). Right words and left words: Electrophysiological evidence for hemispheric differences in meaning processing. Cognitive Brain Research, 8(3), 373–392.

Federmeier, K.D., Wlotko, E.W., & Meyer, A.M. (2008). What’s ‘right’ in language comprehension: Event-related potentials reveal right hemisphere language capabilities. Language and linguistics compass, 2(1), 1–17.

Fedorenko, E., Behr, M. K., & Kanwisher, N. (2011). Functional specificity for high-level linguistic processing in the human brain. Proceedings of the National Academy of Sciences of the United States of America, 108(39), 16428–16433.

Fedorenko, E., Duncan, J., Kanwisher, N. (2013). Broad domain generality in focal regions of frontal and parietal cortex. Proceedings of the National Academy of Sciences of the United States of America, 110, 16616–16621.

Fedorenko, E. & Blank, I. (in press). Broca’s area is not a natural kind.

Fedorenko, E., Hsieh, P.-J., Nieto-Castañón, A., Whitfield-Gabrieli, S., & Kanwisher, N. (2010). New method for fMRI investigations of language: defining ROIs functionally in individual subjects. Journal of Neurophysiology, 104, 1177–1194.

Fedorenko, E., Mineroff, Z., Siegelman, M., & Blank, I. (2018). Word meanings and sentence structure recruit the same set of fronto-temporal regions during comprehension. bioRxiv, 477851.

Fedorenko, E., Nieto-Castanon, A., & Kanwisher, N. (2012). Lexical and syntactic representations in the brain: an fMRI investigation with multi-voxel pattern analyses. Neuropsychologia, 50(4), 499–513.

Fein, D., Humes, M., Kaplan, E., Lucci, D., & Waterhouse, L. (1984). The question of left hemisphere dysfunction in infantile autism. Psychological Bulletin, 95(2), 258–281.

Fisher, S. E., & Scharff, C. (2009). FOXP2 as a molecular window into speech and language. Trends in Genetics, 25(4), 166–177.

Fischl, B., Rajendran, N., Busa, E., Augustinack, J., Hinds, O., Yeo, B.T., … & Zilles, K. (2008). Cortical folding patterns and predicting cytoarchitecture. Cerebral cortex, 18(8), 1973–1980.

Fombonne, E. (2005). Epidemiology of autistic disorder and other pervasive developmental disorders. The Journal of Clinical Psychiatry, 66(10), 3–8.

Friederici, A.D., Brauer, J., & Lohmann, G. (2011). Maturation of the language network: from inter-to intrahemispheric connectivities. PLoS One, 6(6).

Frost, M.A., & Goebel, R. (2012). Measuring structural-functional correspondence: Spatial variability of specialised brain regions after macro-anatomical alignment. NeuroImage, 59(2), 1369–1381.

Gallagher, H.L., Happé, F., Brunswick, N., Fletcher, P.C., Frith, U., & Frith, C.D. (2000). Reading the mind in cartoons and stories: An fMRI study of “theory of mind” in verbal and nonverbal tasks. Neuropsychologia, 38(1), 11–21.

Gaugler, T., Klei, L., Sanders, S.J., Bodea, C.A., Goldberg, A.P., Lee, A.B., … Buxbaum, J.D. (2014). Most genetic risk for autism resides with common variation. Nature Genetics, 46(8), 881–885.

Geschwind, D.H. (2011). Genetics of autism spectrum disorders. Trends in cognitive sciences, 15(9), 409–416.

Geschwind, D.H., Miller, B.L., DeCarli, C., & Carmelli, D. (2002). Heritability of lobar brain volumes in twins supports genetic models of cerebral laterality and handedness. Proceedings of the National Academy of Sciences of the United States of America, 99(5), 3176–3181. https://doi.org/10.1073/pnas.052494999

Gilbert, A.L., Regier, T., Kay, P., & Ivry, R.B. (2006). Whorf hypothesis is supported in the right visual field but not the left. Proceedings of the National Academy of Sciences of the United States of America, 103(2), 489–494.

Gotts, S.J., Jo, H.J., Wallace, G.L., Saad, Z.S., Cox, R.W., & Martin, A. (2013). Two distinct forms of functional lateralization in the human brain. Proceedings of the National Academy of Sciences of the United States of America, 110(36), E3435–E3444.

Grandin, T. (2006). Thinking in pictures: And other reports from my life with autism. Vintage.

Greenberg, D.M., Warrier, V., Allison, C., & Baron-Cohen, S. (2018). Testing the empathizing–systemizing theory of sex differences and the extreme male brain theory of autism in half a million people. Proceedings of the National Academy of Sciences of the United States of America, 115(48), 12152–12157.

Hadjikhani, N., Joseph, R.H., Snyder, J., & Tager-Flusberg, H. (2007). Abnormal activation of the social brain during face perception in autism. Human Brain Mapping, 28(5), 441–449.

Hale, T. S., McCracken, J. T., McGough, J. J., Smalley, S. L., Phillips, J. M., & Zaidel, E. (2005). Impaired linguistic processing and atypical brain laterality in adults with ADHD. Clinical Neuroscience Research, 5(5-6), 255–263.

Hamilton, R.H., Sanders, L., Benson, J., Faseyitan, O., Norise, C., Naeser, M., Martin, P., Coslett, H.B. (2010). Stimulating conversation: enhancement of elicited propositional speech in a patient with chronic non-fluent aphasia following transcranial magnetic stimulation. Brain and Language, 113(1), 45–50.

Harris, G.J., Chabris, C.F., Clark, J., Urban, T., Aharon, I., Steele, S., … Tager-Flusberg, H. (2006). Brain activation during semantic processing in autism spectrum disorders via functional magnetic resonance imaging. Brain and Cognition, 61(1), 54–68.

He, Y., Byrge, L., & Kennedy, D.P. (in press). Nonreplication of functional connectivity differences in autism spectrum disorder across multiple sites and denoising strategies. Human Brain Mapping.

Hedge, C., Powell, G., & Sumner, P. (2018). The reliability paradox: Why robust cognitive tasks do not produce reliable individual differences. Behavior Research Methods, 50(3), 1166–1186.

Herbert, M. R., Harris, G. J., Adrien, K. T., Ziegler, D. A., Makris, N., Kennedy, D. N., … Caviness, V. S. (2002). Abnormal asymmetry in language association cortex in autism. Annals of Neurology, 52(5), 588–596.

Herringshaw, A.J., Ammons, C.J., DeRamus, T.P., & Kana, R.K. (2016). Hemispheric differences in language processing in autism spectrum disorders: A meta-analysis of neuroimaging studies. Autism Research, 9(10), 1046–1057.

Hodapp, A. F., & Gerken, K. C. (1999). Correlations between scores for Peabody picture vocabulary test—III and the Wechsler intelligence scale for children—III. Psychological Reports, 84(3_suppl), 1139–1142.

Holland, S. K., Vannest, J., Mecoli, M., Jacola, L. M., Tillema, J.-M., Karunanayaka, P. R., … Byars, A. W. (2007). Functional MRI of language lateralization during development in children. International Journal of Audiology, 46(9), 533–551.

Huguet, G., Ey, E., & Bourgeron, T. (2013). The Genetic Landscapes of Autism Spectrum Disorders. Annual Review of Genomics and Human Genetics, 14(1), 191–213.

Jenkinson, M. (1999). Measuring Transformation Error by RMS Deviation. FMRIB Technical Report, TR99(MJ1), 1–4.

Joseph, R. (1988). The right cerebral hemisphere: Emotion, music, visual-spatial skills, body-image, dreams, and awareness. Journal of clinical psychology, 44(5), 630–673.

Juch, H., Zimine, I., Seghier, M. L., Lazeyras, F., & Fasel, J. H. D. (2005). Anatomical variability of the lateral frontal lobe surface: Implication for intersubject variability in language neuroimaging. NeuroImage, 24(2), 504–514.

Just, M. A., Cherkassky, V. L., Keller, T. A., & Minshew, N. J. (2004). Cortical activation and synchronization during sentence comprehension in high-functioning autism: Evidence of underconnectivity. Brain, 127(8), 1811–1821.

Kaufman, A. S. (1990). Kaufman brief intelligence test: KBIT. Circle Pines, MN: AGS, American Guidance Service.

Kenworthy, L., Wallace, G. L., Birn, R., Milleville, S. C., Case, L. K., Bandettini, P. A., & Martin, A. (2013). Aberrant neural mediation of verbal fluency in autism spectrum disorders. Brain and cognition, 83(2), 218–226.

Kleinhans, N. M., Müller, R. A., Cohen, D. N., & Courchesne, E. (2008). Atypical functional lateralization of language in autism spectrum disorders. Brain Research, 1221, 115–125.

Knaus, T. A., Silver, A. M., Kennedy, M., Lindgren, K. A., Dominick, K. C., Siegel, J., & Tager-Flusberg, H. (2010). Language laterality in autism spectrum disorder and typical controls: A functional, volumetric, and diffusion tensor MRI study. Brain and Language, 112(2), 113–120.

Knaus, T. A., Silver, A. M., Lindgren, K. A., Hadjikhani, N., & Tager-Flusberg, H. (2008). fMRI activation during a language task in adolescents with ASD. Journal of the International Neuropsychological Society, 14(6), 967.

Koldewyn, K., Yendiki, A., Weigelt, S., Gweon, H., Julian, J., Richardson, H., … Kanwisher, N. (2014). Differences in the right inferior longitudinal fasciculus but no general disruption of white matter tracts in children with autism spectrum disorder. Proceedings of the National Academy of Sciences of the United States of America, 111(5), 1981–1986.

Kriegeskorte, N., Simmons, W. K., Bellgowan, P. S., & Baker, C. I. (2009). Circular analysis in systems neuroscience: the dangers of double dipping. Nature neuroscience, 12(5), 535.

Lai, M.C., Lombardo, M.V., Auyeung, B., Chakrabarti, B., & Baron-Cohen, S. (2015). Sex/gender differences and autism: setting the scene for future research. Journal of the American Academy of Child & Adolescent Psychiatry, 54(1), 11–24.

Landry, O., & Chouinard, P. A. (2016). Why We Should Study the Broader Autism Phenotype in Typically Developing Populations. Journal of Cognition and Development, 17(4), 584–595.

Lee, M. D., & Wagenmakers, E. J. (2013). Bayesian data analysis for cognitive science: A practical course.

Lindell, A. K. (2006). In your right mind: Right hemisphere contributions to language processing and production. Neuropsychology Review, 16(3), 131–148.

Lindell, A. K., & Hudry, K. (2013). Atypicalities in cortical structure, handedness, and functional lateralization for language in autism spectrum disorders. Neuropsychology Review.

Lord, C., Petkova, E., Hus, V., Gan, W., Lu, F., Martin, D. M., … Risi, S. (2012). A multisite study of the clinical diagnosis of different autism spectrum disorders. Archives of General Psychiatry, 69(3), 306–313. h

Lord, C., Rutter, M., DiLavore, P. C., & Risi, S. (1999). Autism Diagnostic Observation Scale-WPS (ADOS-WPS). Los Angeles: Western Psychological Services.

MacDermot, K.D., Bonora, E., Sykes, N., Coupe, A.-M., Lai, C.S.L., Vernes, S.C., … Fisher, S.E. (2005). Identification of FOXP2 Truncation as a Novel Cause of Developmental Speech and Language Deficits. The American Journal of Human Genetics, 76(6), 1074–1080.

Mahowald, K., & Fedorenko, E. (2016). Reliable individual-level neural markers of high-level language processing: A necessary precursor for relating neural variability to behavioral and genetic variability. Neuroimage, 139, 74–93.

Maljaars, J., Noens, I., Scholte, E., & van Berckelaer-Onnes, I. (2012). Language in low-functioning children with autistic disorder: Differences between receptive and expressive skills and concurrent predictors of language. Journal of Autism and Developmental Disorders, 42(10), 2181–2191.

Mason, R. A., Williams, D. L., Kana, R. K., Minshew, N., & Just, M. A. (2008). Theory of Mind disruption and recruitment of the right hemisphere during narrative comprehension in autism. Neuropsychologia, 46(1), 269–280.

McNealy, K., Mazziotta, J. C., & Dapretto, M. (2011). Age and experience shape developmental changes in the neural basis of language-related learning. Developmental science, 14(6), 1261–1282.

Miles, J. H. (2011). Autism spectrum disorders-a genetics review. Genetics in Medicine□: Official Journal of the American College of Medical Genetics, 13(4), 278–294.

Minagawa-Kawai, Y., Van Der Lely, H., Ramus, F., Sato, Y., Mazuka, R., & Dupoux, E. (2011). Optical brain imaging reveals general auditory and language-specific processing in early infant development. Cerebral Cortex, 21(2), 254–261.

Mitchell, R. L. C., & Crow, T. J. (2005). Right hemisphere language functions and schizophrenia: The forgotten hemisphere? Brain.

Müller, R. A., Kleinhans, N., Kemmotsu, N., Pierce, K., & Courchesne, E. (2003). Abnormal variability and distribution of functional maps in autism: An fMRI study of visuomotor learning. American Journal of Psychiatry, 160(10), 1847–1862.

Muhle, R., Trentacoste, S. V., & Rapin, I. (2004). The genetics of autism. Pediatrics, 113(5), e472–e486.

Newbury, D.F., Bonora, E., Lamb, J.A., Fisher, S.E., Lai, C.S., Baird, G., … & Bolton, P.F. (2002). FOXP2 is not a major susceptibility gene for autism or specific language impairment. The American Journal of Human Genetics, 70(5), 1318–1327.

Nielsen, J.A., Zielinski, B.A., Fletcher, P., Alexander, A.L., Lange, N., Bigler, E.D., … Anderson, J.S. (2014). Abnormal lateralization of functional connectivity between language and default mode regions in autism. Molecular Autism, 5(1), 8.

Nieto-Castañon, A., & Fedorenko, E. (2012). Subject-specific functional localizers increase sensitivity and functional resolution of multi-subject analyses. Neuroimage, 63, 1646–1669.

Norman-Haignere, S., Kanwisher, N. G., & McDermott, J. H. (2015). Distinct cortical pathways for music and speech revealed by hypothesis-free voxel decomposition. Neuron, 88(6), 1281–1296.

Oertel-Knöchel, V., & Linden, D. E. J. (2011). Cerebral Asymmetry in Schizophrenia. The Neuroscientist, 17(5), 456–467.

Pardo, C. A., & Eberhart, C. G. (2007). The neurobiology of autism. Brain Pathology, 17(4), 434–447.

Pena, M., Maki, A., Kovac□ić, D., Dehaene-Lambertz, G., Koizumi, H., Bouquet, F., & Mehler, J. (2003). Sounds and silence: an optical topography study of language recognition at birth. Proceedings of the National Academy of Sciences of the United States of America, 100(20), 11702–11705.

Peelle, J. E. (2012). The hemispheric lateralization of speech processing depends on what “speech” is: a hierarchical perspective. Frontiers in human neuroscience, 6, 309.

Pierce, K. (2001). Face processing occurs outside the fusiform ‘face area’ in autism: evidence from functional MRI. Brain, 124(10), 2059–2073.

Piven, J., Palmer, P., Jacobi, D., Childress, D., & Arndt, S. (1997). Broader autism phenotype: Evidence from a family history study of multiple-incidence autism families. American Journal of Psychiatry, 154(2), 185–190.

Poeppel, D. (2014). The neuroanatomic and neurophysiological infrastructure for speech and language. Current Opinion in Neurobiology, 28, 142–149.

Postema, M.C., Van Rooij, D., Anagnostou, E., Arango, C., Auzias, G., Behrmann, M., … & Deruelle, C. (2019). Altered structural brain asymmetry in autism spectrum disorder in a study of 54 datasets. Nature communications, 10(1), 1–12.

Redcay, E., & Courchesne, E. (2008). Deviant Functional Magnetic Resonance Imaging Patterns of Brain Activity to Speech in 2-3-Year-Old Children with Autism Spectrum Disorder. Biological Psychiatry, 64(7), 589–598.

Robinson, E.B., St Pourcain, B., Anttila, V., Kosmicki, J.A., Bulik-Sullivan, B., Grove, J., … Daly, M.J. (2016). Genetic risk for autism spectrum disorders and neuropsychiatric variation in the general population. Nature Genetics, 48(5), 552–555.

Roland, P.E., & Friberg, L. (1985). Localization of cortical areas activated by thinking. Journal of Neurophysiology, 53(5), 1219–1243.

Saxe, R., & Kanwisher, N. (2003). People thinking about thinking people: the role of the temporo-parietal junction in “theory of mind”. Neuroimage, 19(4), 1835–1842.

Saxe, R., & Powell, L.J. (2006). It’s the thought that counts: specific brain regions for one component of theory of mind. Psychological science, 17(8), 692–699.

Seghier, M.L., Lazeyras, F., Pegna, A.J., Annoni, J.M., & Khateb, A. (2008). Group analysis and the subject factor in functional magnetic resonance imaging: Analysis of fifty right-handed healthy subjects in a semantic language task. Human Brain Mapping, 29(4), 461–477.

Schönbrodt, F. D., & Perugini, M. (2013). At what sample size do correlations stabilize?. Journal of Research in Personality, 47(5), 609–612.

Scott-Van Zeeland, A.A., Abrahams, B.S., Alvarez-Retuerto, A.I., Sonnenblick, L.I., Rudie, J.D., Ghahremani, D., … & Bookheimer, S.Y. (2010). Altered functional connectivity in frontal lobe circuits is associated with variation in the autism risk gene CNTNAP2. Science translational medicine, 2(56), 5680.

Szaflarski, J.P., Binder, J.R., Possing, E.T., McKiernan, K.A., Ward, B.D., & Hammeke, T.A. (2002). Language lateralization in left-handed and ambidextrous people: fMRI data. Neurology, 59(2), 238–244.

Sun, T., Patoine, C., Abu-Khalil, A., Visvader, J., Sum, E., Cherry, T. J., … & Walsh, C. A. (2005). Early asymmetry of gene transcription in embryonic human left and right cerebral cortex. Science, 308(5729), 1794–1798.

Tager-Flusberg, H., & Kasari, C. (2013). Minimally verbal school-aged children with autism spectrum disorder: The neglected end of the spectrum. Autism research, 6(6), 468–478.

Tager-Flusberg, H., Paul, R., & Lord, C. (2013). Language and Communication in Autism. In Handbook of Autism and Pervasive Developmental Disorders (pp. 335–364).

Takeuchi, M., Harada, M., Matsuzaki, K., Nishitani, H., & Mori, K. (2004). Difference of signal change by a language task on autistic patients using functional MRI. Journal of Medical Investigation, 51(1–2), 59–62.

Talkowski, M. E., Minikel, E. V., & Gusella, J. F. (2014). Autism spectrum disorder genetics: diverse genes with diverse clinical outcomes. Harvard review of psychiatry, 22(2), 65–75.

Tesink, C. M., Buitelaar, J. K., Petersson, K. M., Van der Gaag, R. J., Kan, C. C., Tendolkar, I., & Hagoort, P. (2009). Neural correlates of pragmatic language comprehension in autism spectrum disorders. Brain, 132(7), 1941–1952.

Thesen, S., Heid, O., Mueller, E., Schad, L.R. (2000). Prospective Acquisition Correction for head motion with image-based tracking for real-time fMRI. Magnetic Resonance in Medicine, 44, 457–465.

Thompson-Schill, S. L., D’Esposito, M., Aguirre, G. K., & Farah, M. J. (1997). Role of left inferior prefrontal cortex in retrieval of semantic knowledge: a reevaluation. Proceedings of the National Academy of Sciences of the United States of America, 94(26), 14792–14797.

Toga, A. W., & Thompson, P. M. (2003). Mapping brain asymmetry. Nature Reviews Neuroscience, 4(1), 37–48.

Tomaiuolo, F., MacDonald, J.D., Caramanos, Z., Posner, G., Chiavaras, M., Evans, A.C., & Petrides, M. (1999). Morphology, morphometry and probability mapping of the pars opercularis of the inferior frontal gyrus: An in vivo MRI analysis. European Journal of Neuroscience, 11(9), 3033–3046.

Turkeltaub, P.E., Coslett, H.B., Thomas, A.L., Faseyitan, O., Benson, J., Norise, C., & Hamilton, R.H. (2012). The right hemisphere is not unitary in its role in aphasia recovery. Cortex, 48(9), 1179–1186.

Tzourio-Mazoyer, N., Landeau, B., Papathanassiou, D., Crivello, F., Etard, O., Delcroix, N.,, et al., (2002). Automated anatomical labeling of activations in SPM using a macroscopic anatomical parcellation of the MNI MRI single-subject brain. Neuroimage,15, 273–289.

Vázquez-Rodríguez, B., Suárez, L. E., Markello, R. D., Shafiei, G., Paquola, C., Hagmann, P., … & Misic, B. (2019). Gradients of structure–function tethering across neocortex. Proceedings of the National Academy of Sciences of the United States of America, 116(42), 21219–21227.

Volkmar, F. R., Lord, C., Bailey, A., Schultz, R. T., & Klin, A. (2004). Autism and pervasive developmental disorders. Journal of Child Psychology and Psychiatry and Allied Disciplines.

Wagenmakers, E. J., Wetzels, R., Borsboom, D., & Van Der Maas, H. L. (2011). Why psychologists must change the way they analyze their data: the case of psi: comment on Bem, Journal of Personality and Social Psychology, 100(3), 426–432

Wang, A. T., Lee, S. S., Sigman, M., & Dapretto, M. (2006). Neural basis of irony comprehension in children with autism: The role of prosody and context. Brain, 129(4), 932–943.

Wehner, D. T., Ahlfors, S. P., & Mody, M. (2007). Effects of phonological contrast on auditory word discrimination in children with and without reading disability: A magnetoencephalography (MEG) study. Neuropsychologia, 45(14), 3251–3262.

Wilkinson, K. M. (1998). Profiles of language and communication skills in autism. Mental Retardation and Developmental Disabilities Research Reviews.

Wing, L. (1988). The continuum of autistic characteristics. In Diagnosis and assessment in autism (pp. 91–110). Springer, Boston, MA.

Yuan, W., Szaflarski, J. P., Schmithorst, V. J., Schapiro, M., Byars, A. W., Strawsburg, R. H., & Holland, S. K. (2006). fMRI shows atypical language lateralization in pediatric epilepsy patients. Epilepsia, 47(3), 593–600.

